# Structural characterization of a novel luciferase-like-monooxygenase from *Pseudomonas meliae* – an in-silico approach

**DOI:** 10.1101/2023.03.27.534437

**Authors:** Mohammad Rayhan, Mohd. Faijanur-Rob Siddiquee, Asif Shahriar, Hossain Ahmed, Aar Rafi Mahmud, Muhammad Shaiful Alam, Muhammad Ramiz Uddin, Mrityunjoy Acharjee, Mst. Sharmin Sultana Shimu, Mohd. Shahir Shamsir, Talha Bin Emran

## Abstract

**Background:** Luciferase is a well-known oxidative enzyme that produces bioluminescence. The *Pseudomonas meliae* is a plant pathogen that causes wood rot on nectarine and peach and possesses a luciferase-like monooxygenase. After activation, it produces bioluminescence, and the pathogen’s bioluminescence is a visual indicator of diseased plants.

**Methods:** The present study aims to model and characterize the luciferase-like monooxygenase protein in *P. meliae* for its similarity to well-established luciferase. In this study, the luciferase-like monooxygenase from *P. meliae* infects chinaberry plants has been modeled first and then studied by comparing it with existing known luciferase. Also, the similarities between uncharacterized luciferase from *P. meliae* and template from *Geobacillus thermodenitrificans* were analyzed to find the novelty of *P. meliae*.

**Results:** The results suggest that the absence of bioluminescence in *P. meliae* could be due to the evolutionary mutation in positions 138 and 311. The active site remains identical except for two amino acids; *P. meliae* Tyr138 instead of His138 and Leu311 instead of His311. Therefore, the *P. meliae* will have a potential future application, and mutation of the residues 138 and 311 can be restored luciferase light-emitting ability.

**Conclusions:** This study will help further improve, activate, and repurpose the luciferase from *P. meliae* as a reporter for gene expression.

## Background

The name of luciferase and luciferin were introduced by scientist Emil Du Bois-Reymond in 1885 (Roura et al., 2013). Later on, in 1940, Dr Green and McElroy extracted and purified luciferase protein. They isolated the enzyme and determined its structure (England et al., 2016). Emil Du Bois-Reymond used cold water to investigate the components required for the click beetle’s bioluminescence, producing luminescence in the laboratory. He named two extracted components, “Luciferine,” which is the enzyme responsible for the chemical reaction, he named luciferase. Marlene Deluca’s cloning of firefly luciferase (FLuc) in *Escherichia coli* directly allows this technique to be widely utilized in many luciferase systems (De Wet et al., 1985). Firefly luciferase catalyzes the oxidative decarboxylation of luciferin, a 6-hydroxybenzothiazole, oxyluciferin in the ATP, Mg^2+^, and O_2_(Berger et al., 2008) (Koncz et al., 1990). Firefly luciferase has been used as reporter gene in living cell and organisms. For instance, the longer-wavelength light emission enhances the animal’s tissue penetration.

Luciferase, specially sourced from firefly (*Photinus pyralis*), has been utilized as a reporter protein in different assay systems, including gene expression, and was applied in high-throughput screening for drug discovery (Inouye, 2010). Using bioluminescence as a visual cue to signal changes has been well established. This bioluminescence happens in nature in different green growths, microscopic organisms, parasites, and some oceanic creatures, such as jellyfish (Thorne et al., 2010). The luciferase gene is extracted and used to visualize various organisms’ gene expressions as a reporter gene. The first luciferase protein was purified from fireflies in the 1940s (Kirkpatrick et al., 2019).

Firefly luciferase has an established application of luciferase-liken in reporting gene regulation and pharmaceutical screening. The *P. meliae*, is a plant pathogenic bacterium that causes wood rot on a nectarine (Fleiss & Sarkisyan, 2019). The presence of luciferase, like monooxygenase by a pathogen. In this case *P. meliae,* creates fireflies intriguing prospect of pathogen’s bioluminescence as a visual indicator of diseased plants. If the pathogen’s protein can be activated when the plant has been infected, its bioluminescent bacterial gall can identify affected plants. In this study, the luciferase suitability, like monooxygenase from *P. meliae* that infects chinaberry plants, is to be first modeled and studied by comparing it with existing known luciferase (Aeini & Taghavi, 2014).

This study aims to examine the luciferase-like monooxygenase gene found in *P. meliae* for its similarity to well-established luciferase enzymes. The novel protein sequence was modeled and compared with known structures using bioinformatics tools. This study will further develop the luciferase from *P. meliae* as a reporter for gene expression.

## Methods

### Selection and retrieval of protein sequence

The selection and retrieval of the luciferase sequence of *P. meliae* are obtained from the Uniprot database. Specific keywords were used, and the description of the retrieved protein sequence was examined for key features of established luciferase enzymes to select the protein sequence correctly. An alignment search was used using the available BLAST tool to explore the sequence similarities. A separate investigation was performed using the retrieved sequence in the protein databank (www.rcsb.org) to confirm the absence of 3D structure (Gasteiger et al., 2003).

### Composition and physicochemical analysis

The retrieved sequence’s primary structure was analyzed using the ProtParam suite of tools (https://web.expasy.org/protparam/). This tool is used to identify the atomic composition, formula, total number of atoms, half-time, estimated amino acid composition, molecular weight, instability index, pI value, hydrophobicity, and other information of the protein model (O’Malley et al., 2012).

### Sequence alignment

Sequence alignment performed to compare the retrieved protein sequence of *P. meliae* with another similar sequence from the UniProt database (https://www.uniprot.org), and UniProt ID (A0A0P9UTV8). Sequence alignment has also been performed using ESPript 3.0 (http://espript.ibcp.fr/ESPript/ESPript/) to identify the conserved region between 2 sequences(Gouet et al., 2003).

### Secondary structure analysis

The secondary structure obtained from the protein sequence is used to identify the beta-sheet, alpha helix, and amino acid sequence. The predicted secondary structure was also focused on a graphical sequence view on the possibility of perpetration of beta-sheet, alpha helix, and turns. The tool used for the secondary prediction structure was CFSSP, which uses the Chou-Fasman algorithm. CFSSP is free via ExPASy server (https://www.biogem.org/tool/chou-fasman/) (Kumar, 2013) and another tool is Network Protein Sequence Analysis (Combet et al., 2000).

### Tertiary Structure Prediction

#### Template selection

The most accurate computational approach for constructing accurate structural model Alphafold is widely used in many biological applications (Jumper et al., 2021). The 3D structure of a query protein helped template the protein’s sequence alignment. Alphafold has been used for the drug discovery process, making the process cheaper and more manageable. This is also used for drug design, analysis of mutations, and binding mechanisms. For the 3D model, the Alphafold (https://alphafold.ebi.ac.uk/) web tool was used. Alphafold is an important technique in structural biology. It has easily determined the gap between known protein sequences (Varadi et al., 2022).

#### Model validation

The protein 3D model built from AlphaFold were evaluated by ERRAT (https://servicesn.mbi.ucla.edu/ERRAT/). ERRAT has been used for evaluation of the model quality and stereochemical properties. It also analyzes the statistics of non-bonded interactions between various atoms (Sumitha et al., 2020). ERRAT gives the statistics of non-bonded interactions between several atom types and plots the error functions assess versus the position of nine-residue sliding windows in a database. As some atoms are provided non-randomly concerning each other in proteins, protein structure can be verified by differentiating between correctly and incorrectly determined regions of protein structures based on characteristic atomic interaction. PROCHECK (https://www.ebi.ac.uk/thornton-srv/software/PROCHECK/) has been used to check a protein structure’s stereochemical quality, creating several PostScript plot analysis (Hameduh et al., 2020). Verify 3D (https://servicesn.mbi.ucla.edu/Verify3D/) has been used for checking the exactness of the modeled 3D structure.

### Structural comparison

The Alpha Fold Protein Structure Database server was used as a template that has high similarity to predicted 3D model of *P. meliae*. The templatlong-chain alkane monooxygenase (LadA) from *Geobacillus thermodenitrificans* (strain NG80-2) is chosen to compare with luciferase model . Both model and template are superimposed by UCSF ChimeraX software. The template 3B9O and the predicted structure A0A0P9UTV8 was compared and visualized using the UCSF ChimeraX software (Pettersen et al., 2021).

## Results

### Selection and retrieval of protein sequence

Sequence alignment comparison, the protein luciferase-like monooxygenase from *P. meliae* was chosen from UniProt ID A0A0P9UTV8 by searching. The sequence has no known tertiary structure in UniProt and subsequent search in protein databank (www.rcsb.org) confirmed that this sequence has no prior structure. The sequence consists of 466 dimer amino acids which is retrieved from Uniprot, and fits the expected length usually found in luciferase protein families.

### Composition and physicochemical analysis

The amino acid composition of both enzymes was calculated by ProtParam.

The chart has been highlighted by color. Model luciferase-like monooxygenase has been highlighted by blue color and suggested template alkane monooxygenase highlighted by orange color. The bar chart (**Figure 1**) shows the highest differences between luciferase-like monooxygenase and alkane monooxygenase are alanine (A) and leucine (L). The model contains 10.1% alanine (A), and the template contains 6.4% alanine (A). On the other hand, alkane monooxygenase contains more lysine (K) (7.0%) compared to luciferase-like monooxygenase (4.5%), and tyrosine (Y) 5.9% for model and 3.4% for template respectively. alanine rich indicate the more hydrophobicity of the model. All the other amino acids are nearly similar between model luciferase like monooxygenase and suggested template alkane monooxygenase.

**Figure 1.**
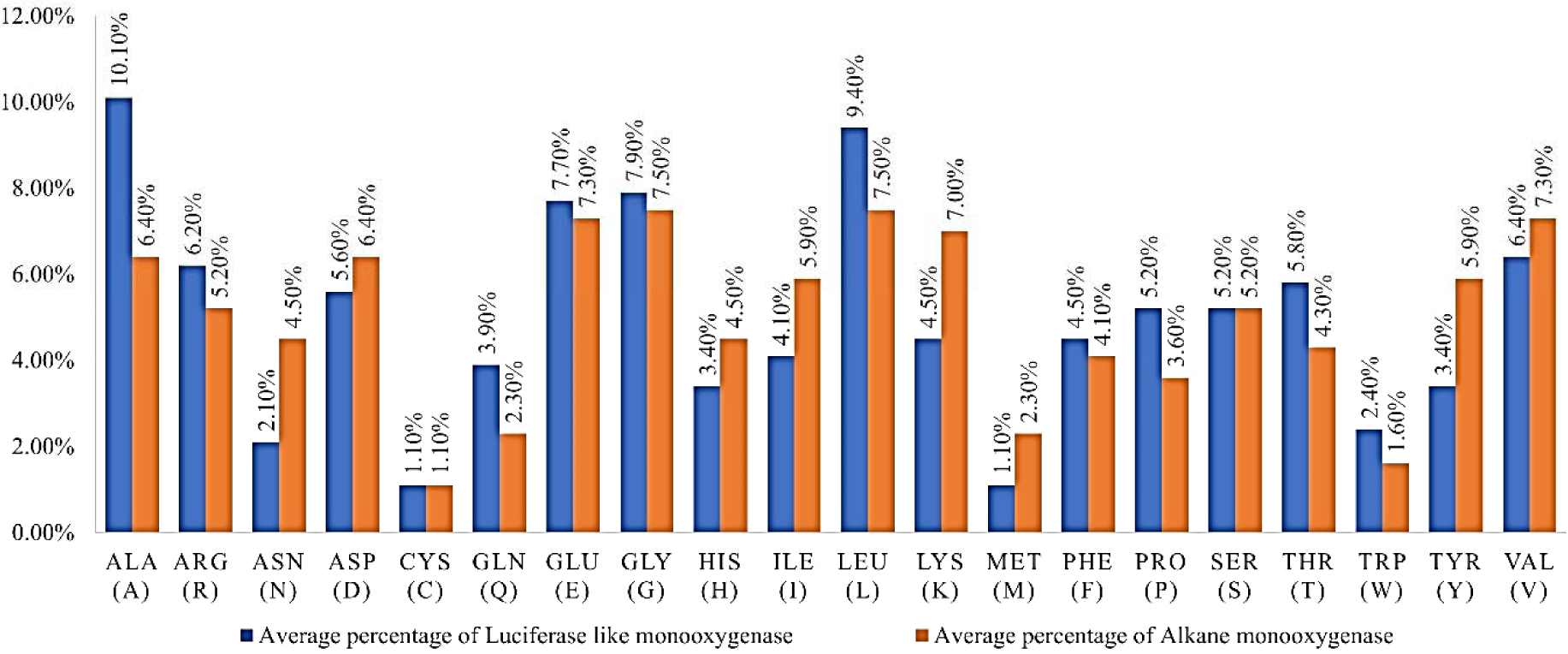
Comparison of the physicochemical characterization in percentages of total residues from luciferase like monooxygenase (blue) and alkane monooxygenase (orange).

The physicochemical characterization of the primary sequence of luciferase-like monooxygenase is composed of 466 amino acid residue, and alkane monooxygenase is composed of 440 amino acid residues. The molecular weight of luciferase-like monooxygenase and alkane monooxygenase is 52.9 and 50.20 kDa respectively. The computed (pI) value of both enzymes is below 7.0. Luciferase-like monooxygenase has a pI value of 5.77, and alkane monooxygenase has 6.38, which means both enzymes are acidic in nature. The study’s finding is consistent with the presence of high acidic residue with different amino acids (Asp and Glu) in luciferase-like monooxygenase and alkane monooxygenase. The similarity in size and acidity suggests that the model may serve similar functions. There was a total 62 negatively charged residue (-R) and 50 positively charged residue (+R) in luciferase-like monooxygenase, and 60 negatively charged residue (-R) and positively charged residue (+54) in alkane monooxygenase respectively. The total number of atoms of luciferase-like monooxygenase is 7,287, and for alkane luciferase, it is 7,035. The total number of atoms luciferase-like monooxygenase is more than the alklane luciferase. Luciferase-like monooxygenase instability index is relatively higher than alkane monooxygenase. The instability index is 38.14 and 29.06, respectively. Grand average of hydropathicity (GRAVY) of luciferase-like monooxygenase is −0.336 and for alkane monooxygenase it is −0.466. The GRAVY value for a protein or a peptide is calculated by adding the hydropathy values of each amino acid residues and then dividing that by the number of residues in the sequence or length of the sequence. Increasing positive score indicates a greater hydrophobicity.

### Sequence alignment

Sequence alignment program for two sequences placed instead use pairwise sequence alignment tools(Chatzou et al., 2016). The program of ESPript (Easy Sequencing in PostScript)(Gouet et al., 2003) allows rapid visualization.The sequence has been aligned to compare model luciferase-like monooxygenase from *P. meliae* with template alkane monooxygenase from *Geobacillus thermodenitrificans* (strain NG80-2). The red marked sequence is similar between template and model, while the white marked sequence is different from each other. The similarities between model luciferase-like-monooxygenase and alkane monooxygenase are 46.36%. The sequence showed alpha helix α (1-14) and beta sheet β (1-13). Sequence alignment has been performed between the model luciferase-like monooxygenase and the template alkane monooxygenase shown in **Figure 2**.

**Figure 2.**
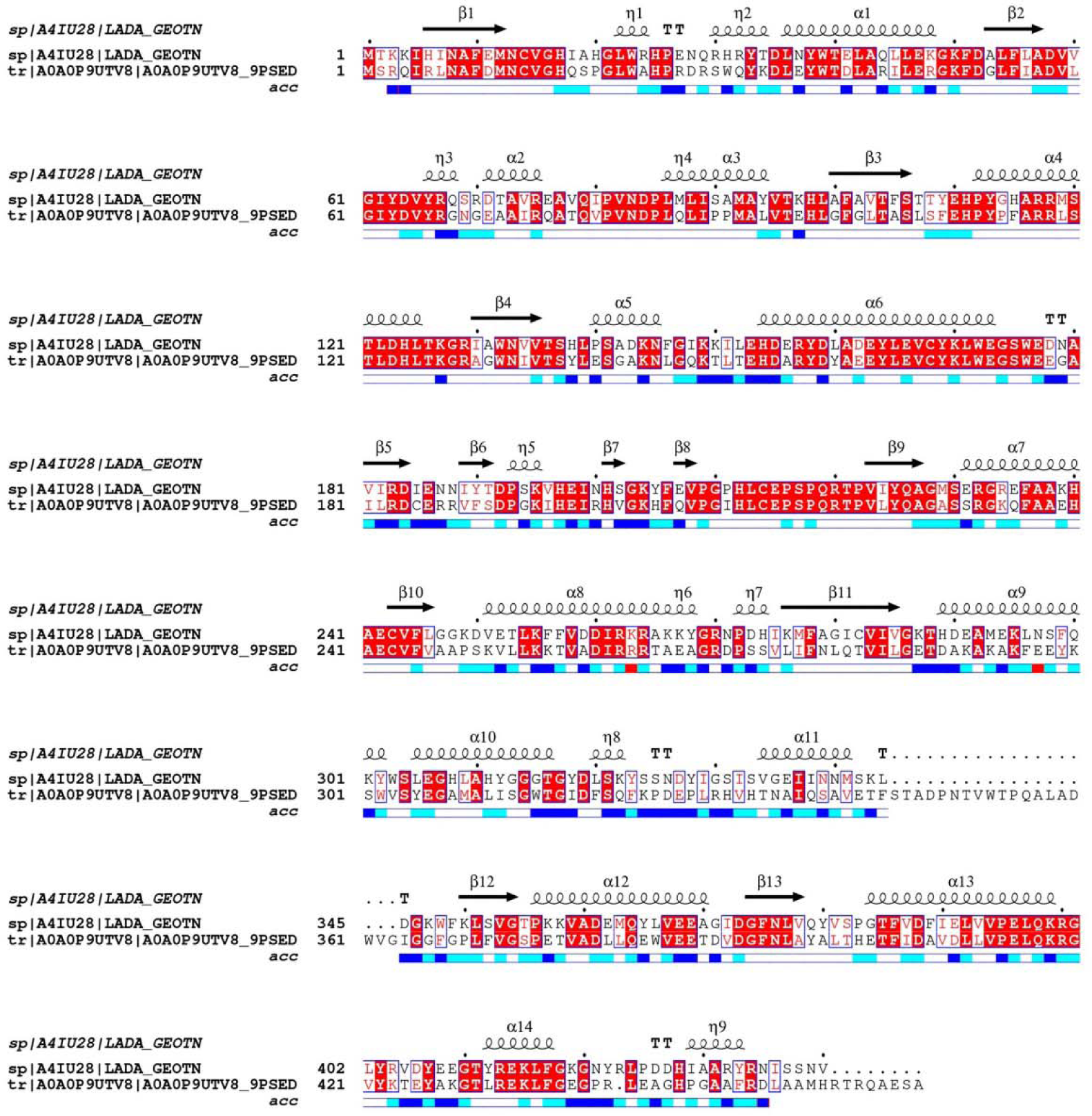
The multiple sequence alignment result between alkane monooxygenase *Geobacillus thermodenitrificans* (strain NG80-2) (top) and luciferase-like monooxygenase gene found in *P. meliae* (bottom). The red bands denote identical residues between the two sequences. The black boxes highlighted the active site of the template and predicted active site of the model.

### Secondary structure analysis

Secondary structure is formed by hydrogen bonds between the amino hydrogen and carboxyl oxygen atoms in the peptide backbone. Secondary structure analysis of protein includes regular secondary structure (helices and strands), which are the main components. Loops are non-regular structure. Both regular and non-regular structures are used for interfacing.

Comparison of the secondary structure using GOR IV (Kouza et al., 2017) showed that the percentage of helices, strand, and coil are similar. The alpha helix presence was dominant in the template (41.85%) and model (35.45%). The number of amino acids is higher in the model. The template has a higher percentage of strands (24.77%) compared to the model (14.38%). The percentage of coil region in template alkane monooxygenase is lower than model luciferase-like monooxygenase. The percentage of the coil is 43.78% for the model and 39.77% for the template. The comparison of model and template has highlighted the similarities in the percentage of alpha helices, beta-sheet, and random coil. According to the percentage of similarities, the random coil is more stable than alpha helices and beta-sheet (**Supplementary Figures S1 and S2**).

### Template Selection

The protein sequence submitted to Alpha fold Protein structure database (Jumper et al., 2021)resulted in the selection of alkane monooxygenase from *Geobacillus thermodenitrificans* (strain NG80-2, PDB ID: 3B9O) and obtained the best score 52.86% with a high percentage of identity and low e-value 2.2e-56 (**Table 1**).

**Table 1.**
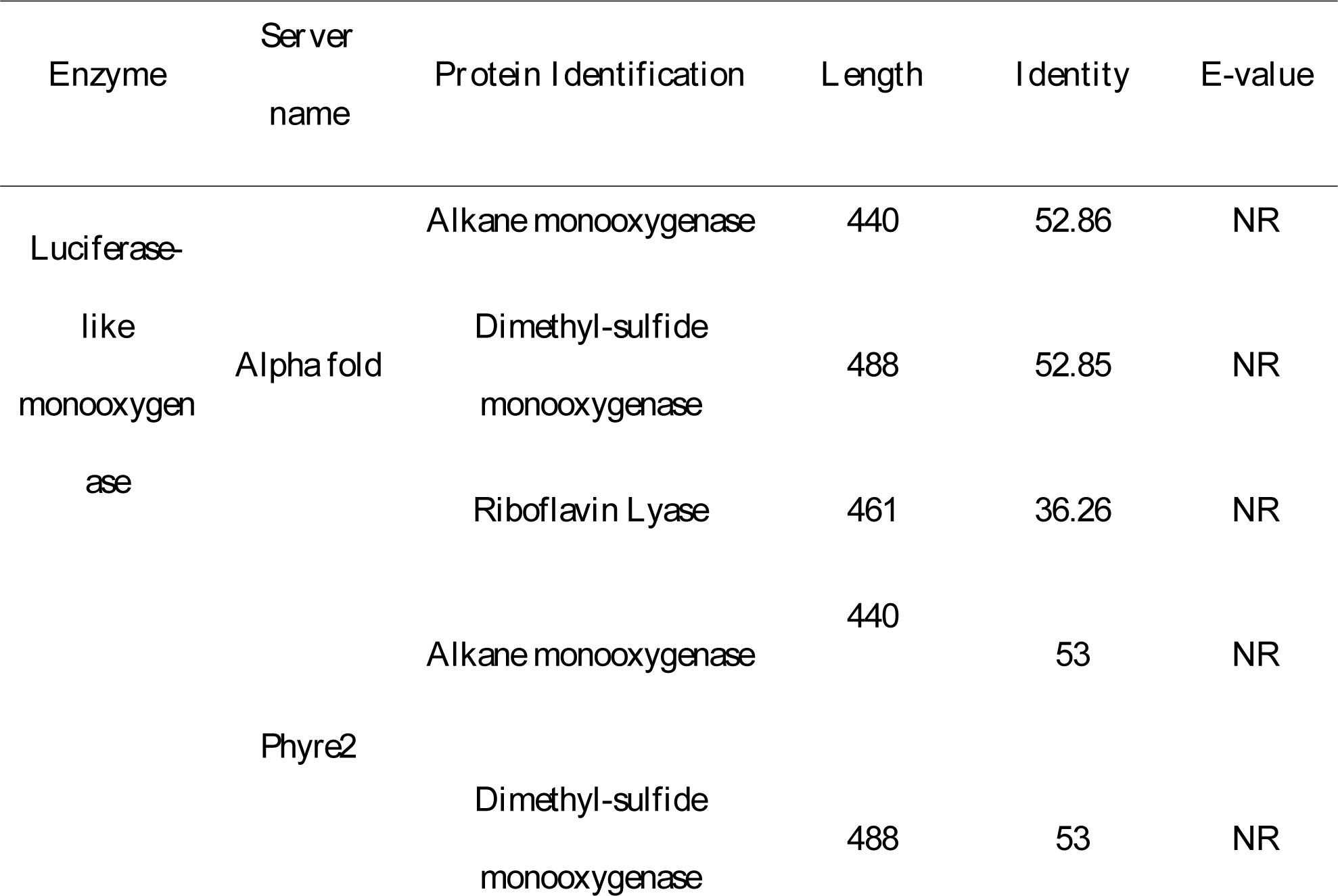
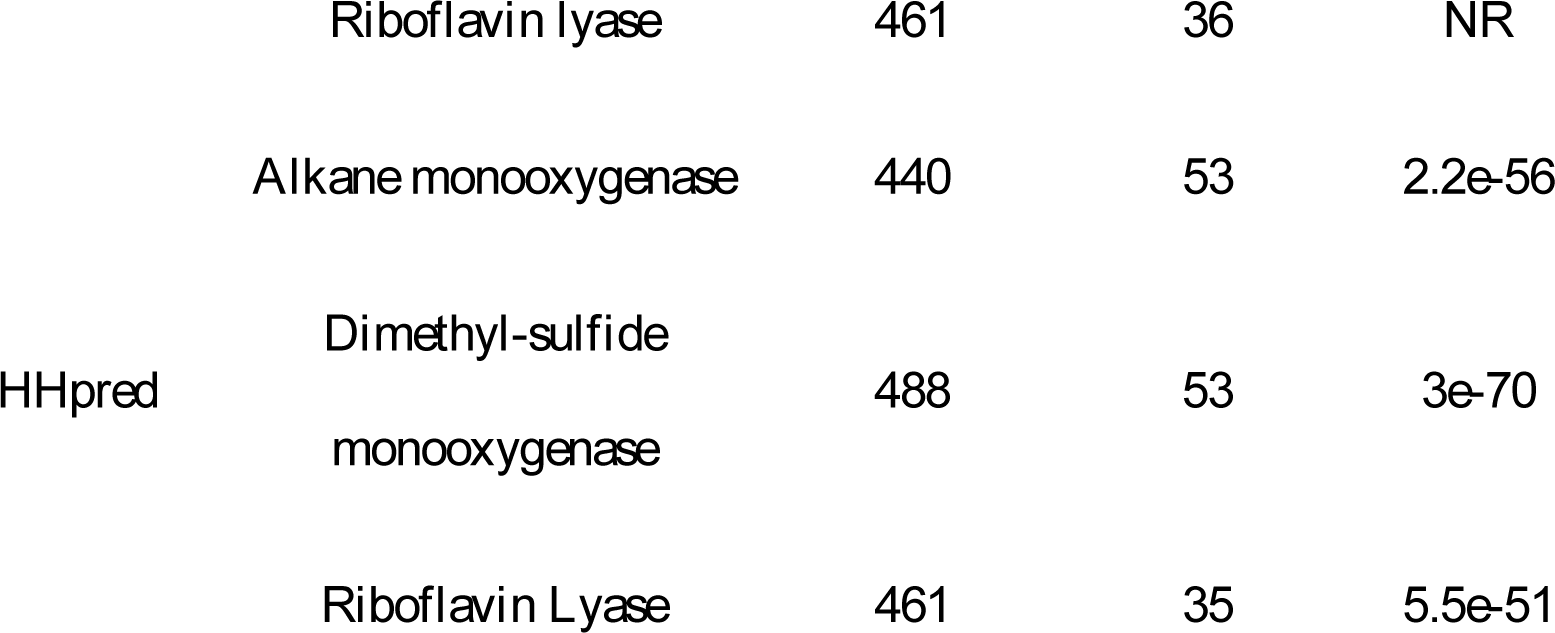
Selected a template for modelling by Accession number, protein name, and organism

Result of the Alpha fold presented in **Supplementary Figure S3**, which shows 46.36% similar identity to the template long-chain alkane monooxygenase (LadA) in thermophilic *bacillus spp* (PDB ID 3B9O).

### Tertiary Structure Prediction

Three-dimensional structure of luciferase-like monooxygenase from *P. meliae* (A0A0P9UTV8) as modelled by using Alpha fold software shown at (**Supplementary Figure S3**). The total chain of the structure started from Methionine (01) and ended of the structure alanine (466)

### Structure validation

#### ERRAT2

The ERRAT software results show a good high-resolution structure, which generally produces overall quality factor, valuing around 95. The overall quality factor of predicted structure luciferase-like monooxygenase from ERRAT2 analysis showed a lower value of 93.231. On the error axis, two lines are drawn to indicate the confidence with which it is possible to reject regions that exceed that error values. Expressed as the percentage of the protein for which is the calculated value falls below the 95% rejection limit. Good high-resolution structure generally produces values around 95% or higher. For lower resolutions (2.5 to 3A) the average overall quality Factor is around 91% (**Figure 3**).

**Figure 3.**
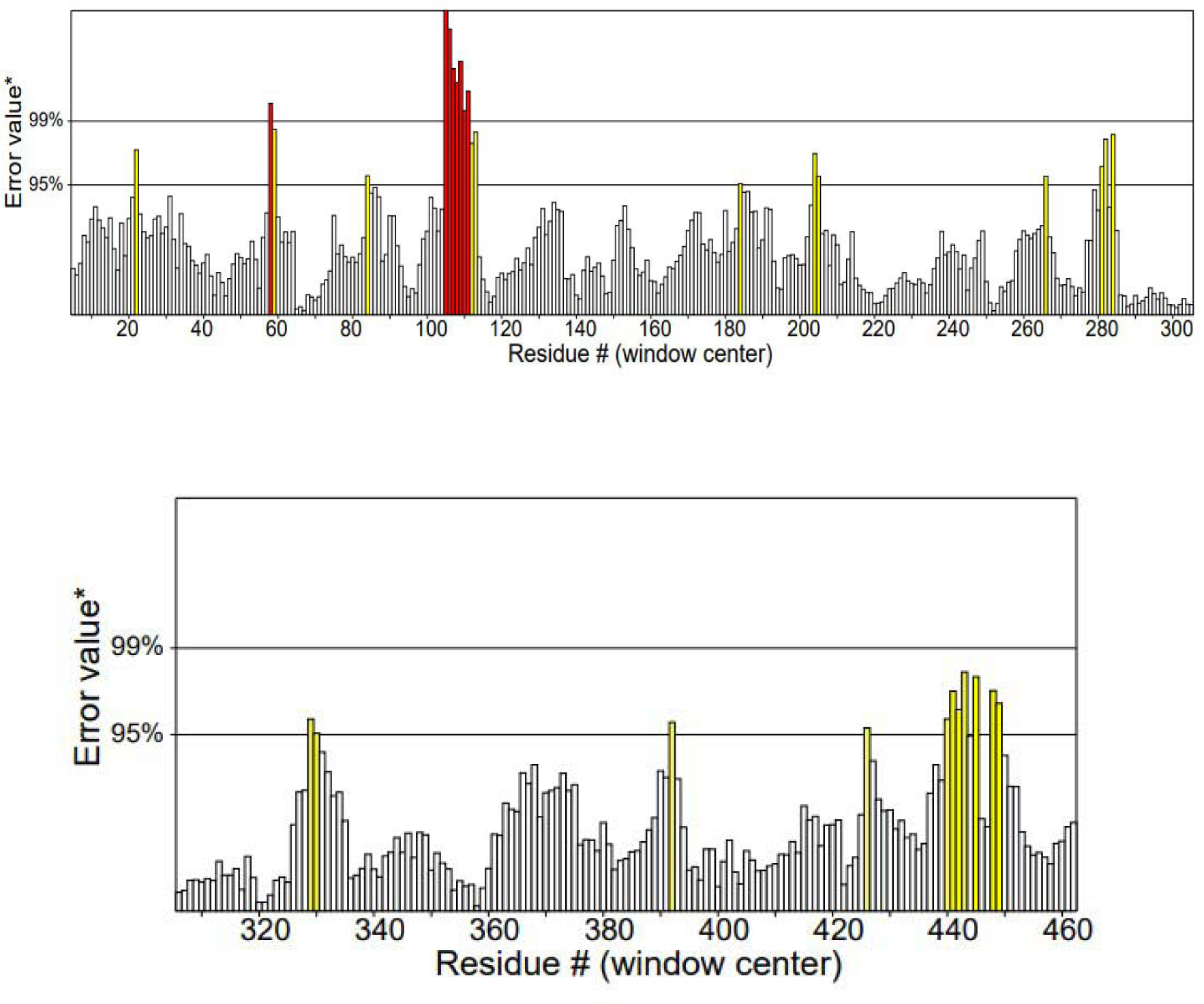
Structure validation using ERRAT2. Two lines are drawn to indicate the possibilities to reject regions that exceed that error value.

#### PROCHECK

The Ramachandran plot (**Figure 4**) shows the validation of the prediction model of structure luciferase-like monooxygenase protein from Alphafold template, as shown below. Ramachandran plot has been shown the different types of areas is that, white areas correspond to conformations where atoms in the polypeptide come closer than the sum of their van der Waals radi. The statistical values, The plot score 92.8%. 5% labelled residues of all Ramachandran’s (out of 464), 2 labelled residues of Chi1-Chi2 plots (out of 272). These regions are sterically disallowed for all amino acids except glycine which is unique in that it lacks a side chain. The red regions correspond to conformations where there are no steric clashes, these are the allowed regions namely the alpha-helical and beta-sheet conformations. The yellow areas show the allowed regions if slightly shorter van der Waals radi are used in the calculation, the atoms are allowed to come a little closer together (**Table 2**).

**Figure 4.**
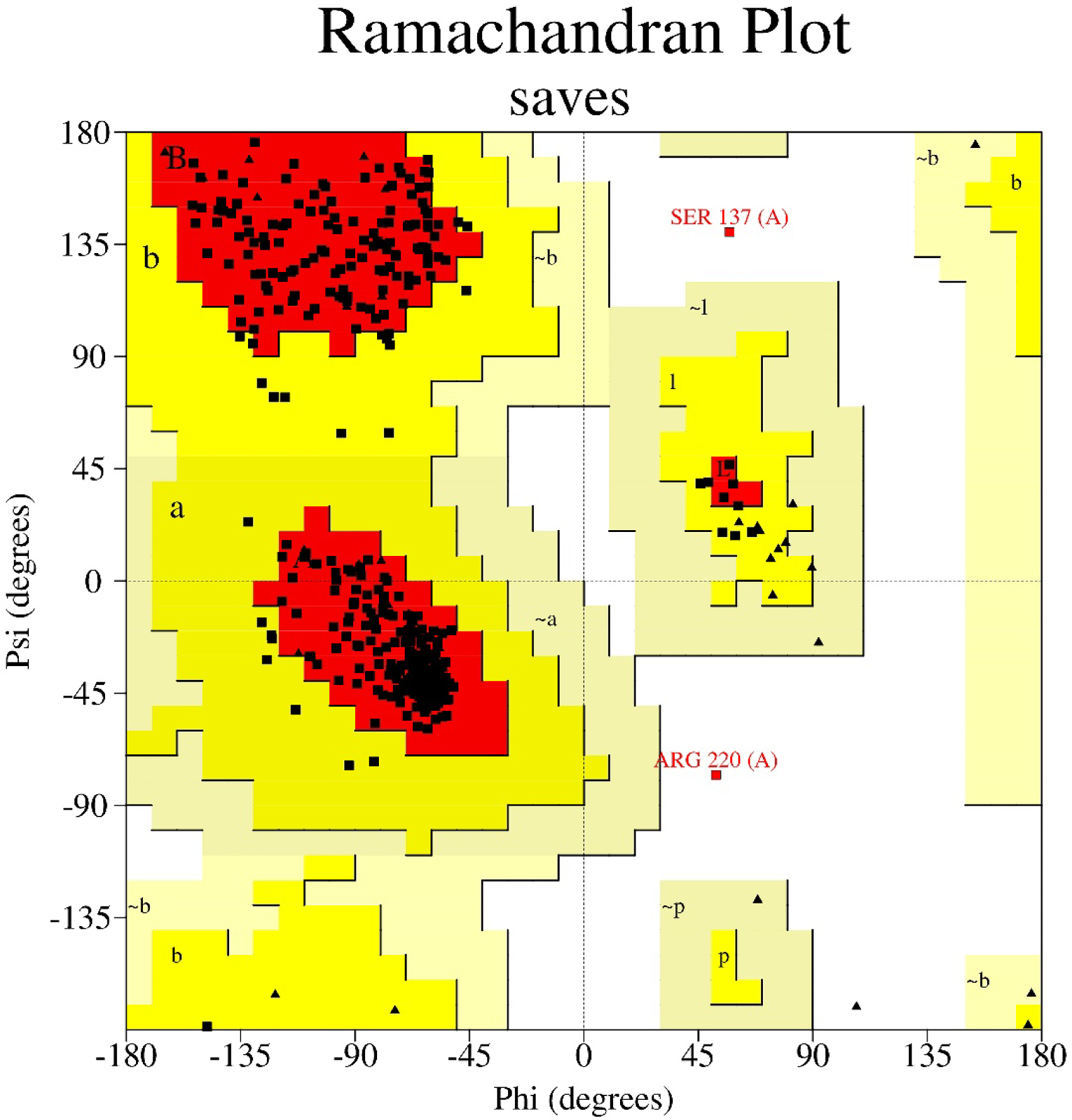
Model validation using Ramachandran plot to show the background of phi-psi probabilities.

**Table 2.** Ramachandran plot validation percentage.

| Evaluation of residues | Score |
| --- | --- |
| Residues in most favoured regions [A,B,L] | (374) 92.8% |
| Residues in additional allowed regions [a,b,l,p] | (27) 6.7% |
| Residues in generously allowed regions [~a,~b,~l,~p] | (0) 0.0% |
| Residues in disallowed regions | (2) 0.5% |
| Number of non-glycine and non-proline residues | (403) 100.0% |
| Number of end-residues (excl. Gly and Pro) | 2 |
| Number of glycine residues (shown as triangles) | 37 |
| Number of proline residues | 24 |
| Total number of residues | 466 |

#### VERIFY 3D

Verify 3D program determines the compatibility of the atomic coordinates of the 3D model with its own amino acid sequence (1D). Three-dimensional profiles computed from correct protein structures match their sequences with high scores. The model structure is passed from validation if at least 80% of the amino acids have scored more excellent or equal to 0.2 in the 3D/1D profile. The quality of the 3D Alphafold structure of the thermophile luciferase protein from Verify3D is as shown below at (**Figure 5**). Around 97.21% of the residues had an average 3D-1D score of ≥ 0.2.

**Figure 5.**
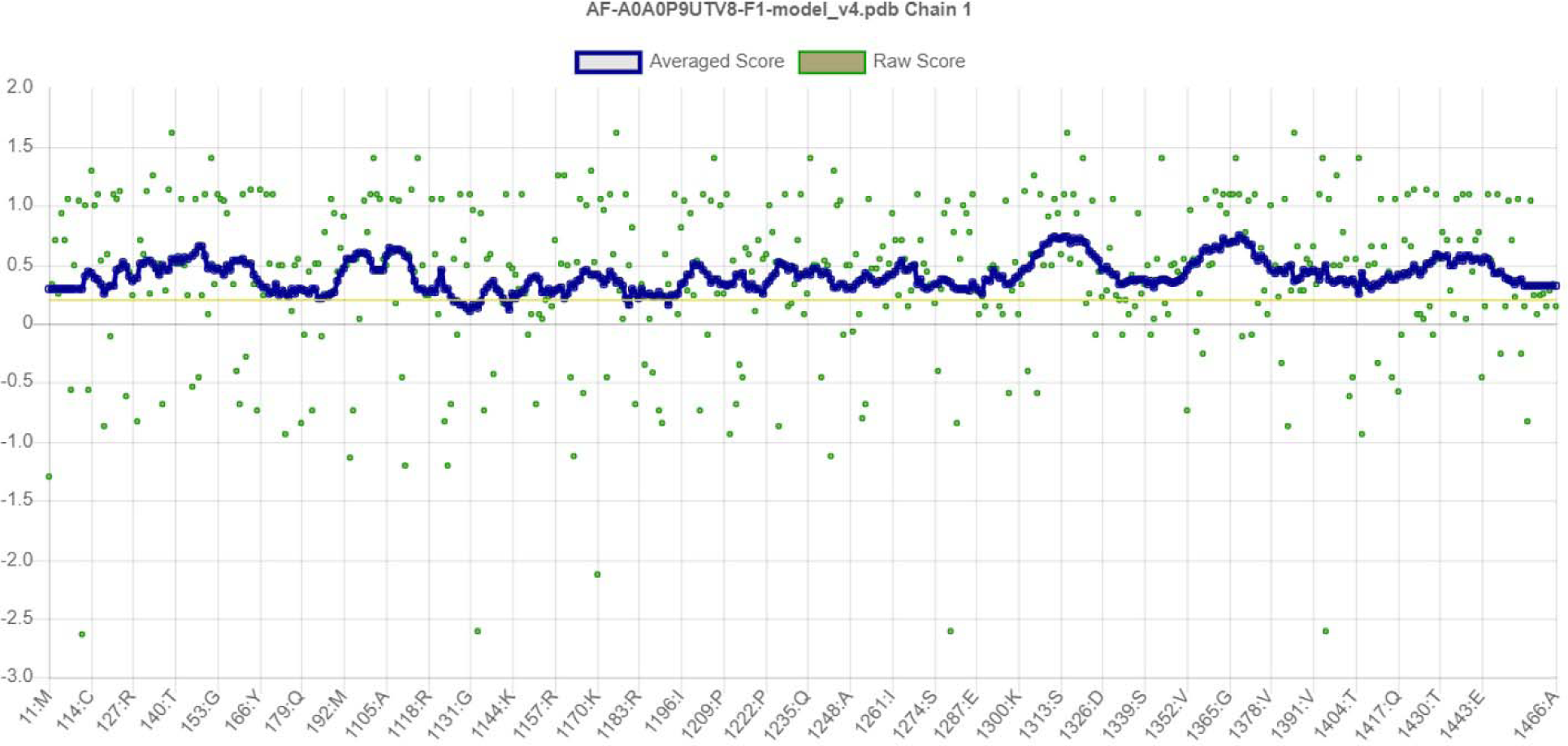
The compatibility of an atomic model (3D) with its own amino acid sequence (1D) by assigning a structural class-based Model validation using Verify 3D.

### Structural comparison

The overall structure and orientation of the modelled structure fits the shape and requirements of a monooxygenase. The only major difference between structure and reference template is the size (26 amino acid length difference). The structure homodimer is composed of two identical subunits. Each subunit contains 466 amino acid residues. The reference template is homodimer composed of 440 amino acid residues. In the **Supplementary Figure 4**, has been shown the surface of the model structure luciferase-like monooxygenase and template alkane monooxygenase.

### Active site prediction

The result showed that the similarity of the model protein luciferase-like monooxygenase with template 3B9O is 46.36%. Since there is no previous research showing the residues’ location that forms the active site, the template 3B9O has been used as a reference to narrow down the possible location of similar amino acids that would come from the active site on model luciferase-like monooxygenase. There are eight important amino acids in the active site, as reported on the Uniprot database for the template 3B9O. Using this as a reference point, all eight amino acids were aligned with the model luciferase-like monooxygenase protein and highlighted. The superimposition of the two structures has been highlighted by using different colors **(Supplementary Figure S5)**. The template alkane monooxygenase and model luciferase-like-monooxygenase both have similarities of amino acids, which has been highlighted by green color and the difference of amino acids has been highlighted by red color. The active site of the template has been highlighted by blue color (**Figure 6**), and the probable active site of the model’s has also been highlighted by blue color (**Figure 7**). The result revealed that the proximity of the 3B9O residues responsible for forming the active site is located near the modelled luciferase residues, even though in the sequence, the active sites of both proteins are located at a distance from each other along the protein sequence. Highlighting the 3D superimposed structure location revealed that both active sites exist at a similar location within the conformational structure of luciferase.

**Figure 6.**
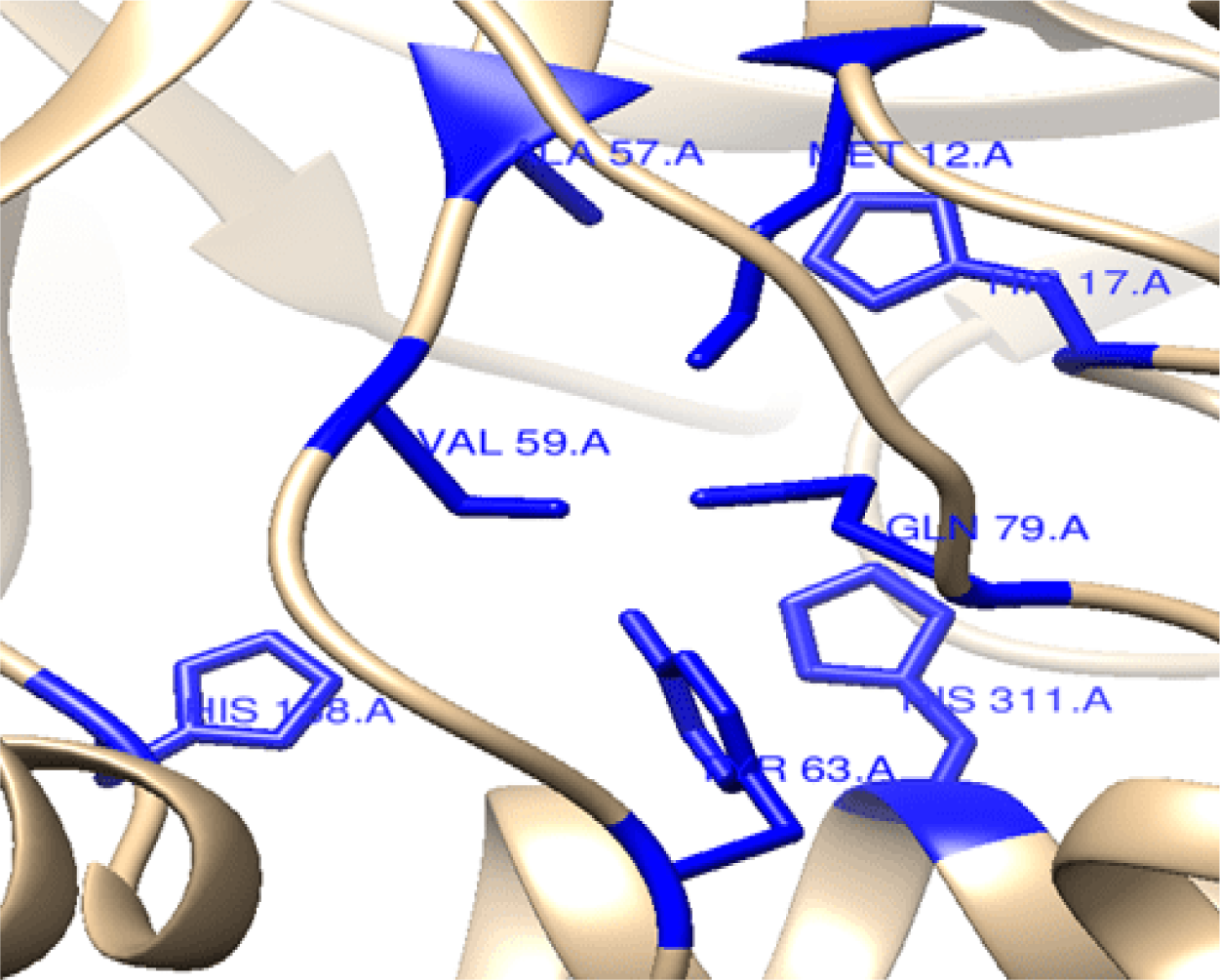
Active site residues of template alkane monooxygenase.

**Figure 7.**
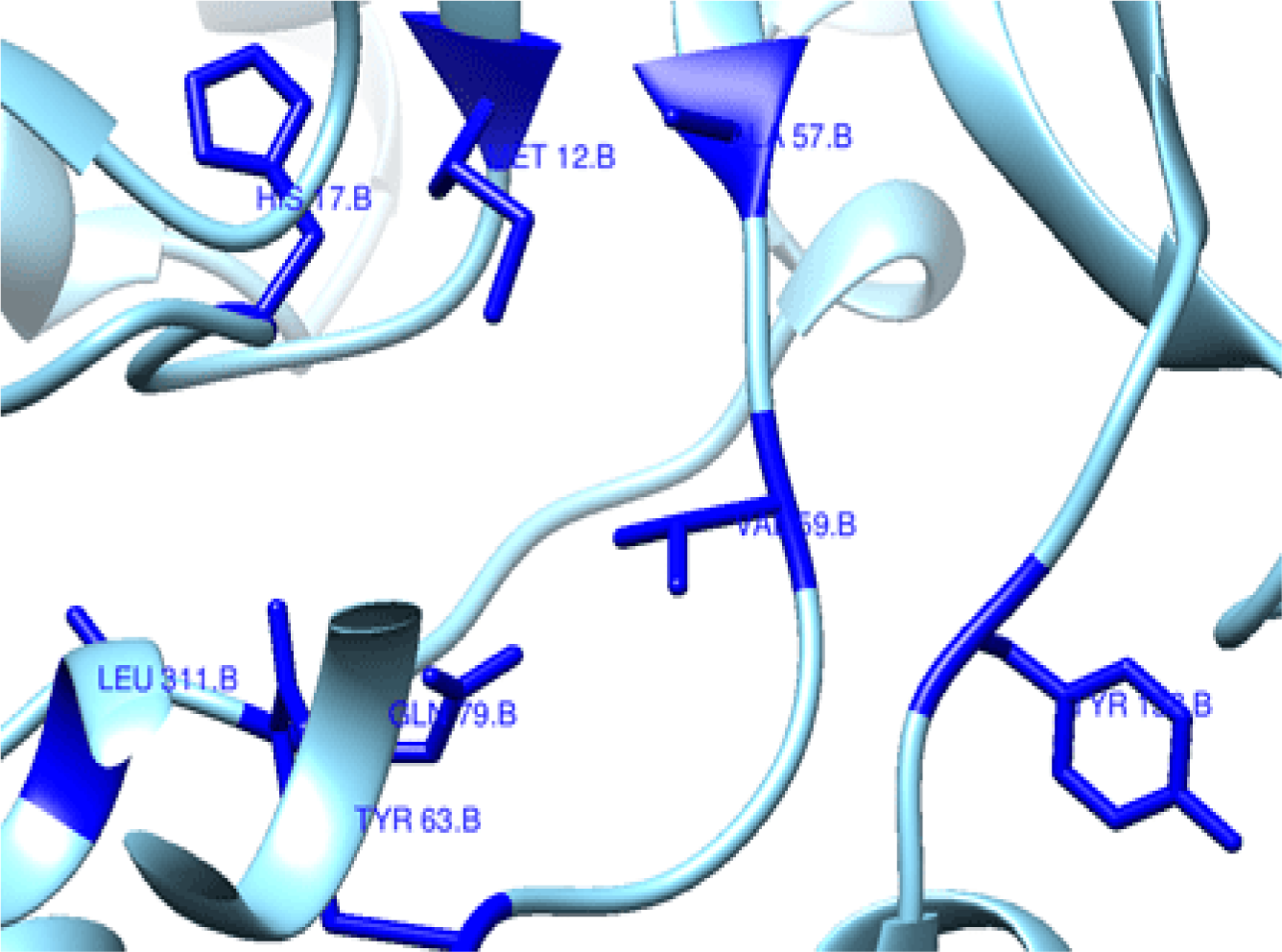
Active site residues of model luciferase-like-monooxygenase.

The figure shows the specific position of the active site of template alkane monooxygenase and the predicted active site of luciferase-like-monooxygenase. At the position of 12, both amino acids are identical to residue methionine. At the 17, both residues are histidine, and at the position of 59, the model and template’s amino acids are identical with Valine. At position 63, the amino acids of template and model are same (Tyrosine). At position 79, template and model are the same amino acids glutamine. At position 138, the template amino acid is histidine and model amino acid is tyrosine. At the position of 311, the template amino acid is histidine, and at the model’s position of 311, the amino acid is leucine. These are only two differences that were discovered between template active site amino acid’s position and model active site amino acid’s position.

The surface will find the whole molecule rather than individual residues. The surface adds an outer picture of molecule, stretched around an imaginary Van der Waals surface underneath. That means only outside of molecule is covered and underlying molecular representation. The surface image has been highlighted for visualize active site of the model and template (**Supplementary Figure S6**). The surface image of active site of template alkane monooxygenase from *Geobacillus thermodenitrificans* (strain NG80-2) (**Supplementary Figure S7**) and predicted active site of luciferase-like-monooxygenase from *P. meliae* (**Supplementary Figure S8**).

### Overall structure analysis

The structural elucidation of the template and model has been highlighted by a different color. The beta-strands are highlighted by blue color, the loops are highlighted by grey color and a shade of light brown highlights helices (**Supplementary Figure S9**).

#### Strands

Beta strand is an important secondary structure element which consists of several beta-strands— stretched segments of the polypeptide chain combined together by the network of hydrogen bonds between adjacent strands. Beta-strand is an important mode of protein-protein interaction(Siepen et al., 2009). The structural difference between template and model strands is the amount of protein. Here, the number of strands in the template structure is 24.77%. The number of strands in the model structure is 14.38% (**Supplementary Figure S10**).

Strands superimpose between model luciferase-like monooxygenase (**Supplementary Figure S11**) and template alkane monooxygenase. The colors have highlighted similarities and differences between model and template. The blue color has highlighted the similarities between model and template, and the red color has highlighted the differences between model and template. The red color at the template and model has specific amino acids, belonging to the same group. The 7th and 224th model positions have Leucine (L) and template have Isoleucine (I), which are the same alkyl group. On the other hand, at 56^th^ position, the model’s amino acid is Isoleucine (I), and the template’s amino acid is Leucine (L), which are the same group. At the 11^th^ position, the model amino acids are aspartic acid (D) and template amino acids are glutamic acid (E). Both are different amino acids, but both amino acids group are same (carboxylic acid group). At the 103^th^ and 285^th^ position, the model’s amino acids are Leucine (L) and template amino acids are Valine (V). Both are not similar but they are from the same alkyl group. But for 246^th^ position, it is reversed. The template’s amino acid is Leucine (L) and model’s amino acid is Valine (V), belonging to the same alkyl group. At the 130^th^ position, the template’s amino acid is Isoleucine (I) and the model’s amino acid Alanine (A), the same alkyl group. The template and model’s amino acids are Valine (V) and Isoleucine (I), but both amino acids belong to the same alkyl group. And finally, at the 397^th^ position, Alanine (A) for model and Valine(V) for template belongs to the same alkyl group(Patel et al., 2021).

#### Loops

Loop regions are one of the most valuable regions in Alpha-fold prediction regions. The coil is a flexible segment of the contiguous polypeptide chain that connects two secondary structures in a protein. The loop region plays critical roles in protein functions, like catalytic sites of enzymes, contributing molecular recognition, and participating in ligand binding sites(Li, 2013). The loop region makes protein function and leads to binding the unaligned region in a sequence alignment between alpha-helix and beta-sheet. The difference between template and structure of loops are already given at (**Supplementary Figure S12**). The template structure composition contains 35.45% of coil or loops. The model structure has 41.85% coils (**Supplementary Figure S13**).

The loops have been highlighted by color. The blue color highlights the similarities between template and model. The red color highlights the differences, but some amino acids are from the same group. For example, Valine (V) and Leucine (L), which are from the same alkyl group, are 40, 400 and 421. Isoleucine (I) and Valine (V) are different amino acids, but they belong to the same alkyl group. They are at the position of 80, 181, 189, 196, 275, and 391. Serine (S) and Threonine (T) are not similar, but they are from the same hydroxyl group and are 108, 191, 374, and 401. At 151 and 182, the template and model have Isoleucine (I) and Leucine (L) from the same alkyl group. There are two different amino acids in the carboxylic acid at 178^th^ and 425^th^, which have Aspartic acid (D) and Glutamic acid (E). At the position of 340, the model has Alanine (A) and the template have Isoleucine (I), but they are from same alkyl group. They have been shown in the same group because of their polarity. Both Alanine and Isoleucine are non-polar and are part of aliphatic amino acid in nature.

#### Helices

Helices are an essential type of secondary structure element found in protein(Fodje & Al-Karadaghi, 2002). Helices play a main role in genome maintenance, how the enzyme changes conformations, and transitions between different conformational states, regulating nucleic acid and reshaping structure(Ma et al., 2018). The number of helices in the template structure is 39.77%, compared to 43.78% helices in the model (**Supplementary Figure S14**).

The color has highlighted similarities and differences between the model template. The blue color has highlighted the similarities between model and template, and the red color has highlighted the differences between model and template. But the red color has some amino acids which belong to the same group. At the 41^th^ position, the model amino acid is aspartic acid (D) and template amino acid is glutamic acid (E), which are different, but from the same group. At the 71^th^ position, model’s amino acid is glutamic acid (E), and the template’s amino acid is aspartic acid (D). At the 74^th^ and 406^th^ position, the model has isoleucine (I) and template have valine (V), and at 409^th^ position, the amino acids are reversed. The template has isoleucine (I) and model has valine (V), but they belong to the same alkyl group. There are two different amino acids at 89, 318, 358, and 364, but they are from the same alkyl group, which also includes isoleucine (I) and valine (V). And the final similar group of amino acid includes valine (V) and leucine (L), at the position of 412 has highlighted given at (**Supplementary Figure S15**).

### Binding site

In proteins, binding sites are small tertiary structure pockets where the ligands bind to it using weak forces (non-covalent bonding)(Patel et al., 2019). The binding sites of protein play an important role in a wide range of applications, including molecular docking, drug design, structure identification, and comparison of functional sites(Guo & Wang, 2012). One of the receptor’s most fundamental properties is the set of amino acids available for interactions with ligands(Khazanov & Carlson, 2013). The binding site of predicted model luciferase-like monooxygenase from P. meliae has given the Uniprot ID. The binding side’s position Asp58 and Thr104 (**Figure 8A**). The template alkane monooxygenase from *Geobacillus thermodenitrificans (strain NG80-2)* has given the binding side at 58, 104, 158 245 (**Figure 8B**).

**Figure 8.** **(A)** Binding residues of model luciferase-like monooxygenase are of 104 and 58 with ligand. **(B)** Binding residues of template alkane monooxygenase are 104, 58, 158, and 245.

## Discussion

In this study, the sequence of luciferase-like monooxygenase was retrieved from UniProt (Pettersen et al., 2021)(Organism: *P. meliae*, ID A0A0P9UTV8) as there was no known structure in protein databases. The homology and its analysis revealed some important features. In physicochemical comparison, we found maximum differences between luciferase-like monooxygenase and alkane monooxygenase in the percentage of alanine (A) and leucine (L), that is, 10.1% and 6.4%, respectively, while other amino acids were similar. Alanine indicates the more hydrophobicity of the model(Lefèvre et al., 1997). The computed (pI) value of both enzymes is below 7.0. Luciferase-like monooxygenase has a pI value of 5.77, and alkane monooxygenase has 6.38, which means both enzymes are acidic in nature. The hydrophobicity nature also supports by the GRAVY value. Here, the GRAVY value of luciferase-like monooxygenase is −0.336, and for alkane monooxygenase, it is −0.466. An increasing positive score indicates a greater hydrophobicity(Lee et al., 2014). The similarity in size and acidity suggests that the model may serve similar functions. In the instability index, luciferase-like monooxygenase has a relatively higher score than alkane monooxygenase, 38.14 and 29.06, respectively. A less than 40 instability index score indicates greater stability [33].

The active sites of both proteins are located at a distance from each other along the protein sequence. Highlighting the 3D superimposed structure location revealed that both active sites exist at a similar location within the conformational structure of luciferase. In both enzymes, all active residues were similar except positions 138 and 311. At position 138, the template amino acid has histidine, and the model amino acid has tyrosine. At the position of 311, the template amino acid has histidine, and at the model’s position of 311, the amino acid is leucine. The findings indicate that the lack of bioluminescence in *P. meliae* of luciferase-like-monooxygenase is due to an evolutionary mutation of an amino acid at positions 138 and 311. Pseudomonas fluorescens, for example, has been shown to react with light and emit luminescence when exposed to UV light. By measuring light emission, bioluminescent *P. aeruginosa* may be used to determine the antimicrobial efficacy of wound dressings. As a result, if the residues 138 and 311 are mutated to recover luciferase light-emitting capacity through future studies, the *P. meliae* could have a few possible applications in the future. There is still space for luciferase-like-monooxygenase from *P. meliae* to be improved, activated, and repurposed as a disease marker. For potential biological applications, such as living cells and living tissue, fluorescence may be used to detect specific components of complex biomolecular assemblies.

The results showed that the data and subsequent model of the luciferase-like monooxygenase from *P. meliae* showed that the plant pathogen’s protein possesses major characteristics of light-emitting luciferase. The overall structure, protein composition, and profile such as acidity suggest that the protein has not evolved significantly different from all other protein in the light-emitting luciferase family. The active site remains identical but except two amino acids; the model from *P. meliae* Tyr138 instead of His138 position in the template, and Leu311 instead of His311. The model tyrosine belongs to hydroxyl and template Histidine belongs to the amino group. Both amino acids (Leucine and Histidine) functional groups shown L-amino group and carboxylic group under certain biological conditions. All the other data on various properties of luciferase-like monooxygenase protein and its comparison between template alkane monooxygenase and model luciferase-like monooxygenase primary structure characteristics, similarities of amino acids sequences, binding sites and predicted active sites of the model are almost similar. Both structures have similar key characteristics, such as high amino acid residue present in the primary protein structure. Both of them have almost same negative and positive charge, which are (-R) 62, (+R) 50 and (-R) 60, (+R) 54 respectively.

## Conclusions

The results suggest that the absence of bioluminescence in *P. meliae* of luciferase-like-monooxygenase could not be emitted light due to the lacking of mutation of amino acid in the position of 138 and 311. Therefore, the *P. meliae* may have a few potential applications in the future, should be mutated the remaining residues 138 and 311 to explore luciferase light-emitting ability through future research. After that, there will be open a room for further improvement, activation, and repurposing of luciferase-like-monooxygenase from *P. meliae* to use it as a disease marker. The *Pseudomonas* genera have been shown to react with light, such as *P. fluorescens* that emit luminescence under UV light. Bioluminescent *P. aeruginosa* can be used to evaluate the antimicrobial efficacy by releasing light emission. The fluorescence could be used to detect particular components of complex biomolecular assemblies for future biological applications, e.g., living cells and living tissue.

## Acknowledgements

Not applicable.

## Authors’ contributions

Mohammad Rayhan and Mohd. Faijanur-Rob Siddiquee conceived the study, designed the experiments, and wrote the manuscript. Mohammad Rayhan, Mohd. Faijanur-Rob Siddiquee, Asif Shahriar, Muhammad Shaiful Alam, Hossain Ahmed, Aar Rafi Mahmud, Mst. Sharmin Sultana Shimu and Mohd. Shahir Shamsir performed the dry-lab experiments. Muhammad Ramiz Uddin and Mrityunjoy Acharjee were performed the English Editing and Proofreading. Mohd. Shahir Shamsir designed and planned the studies, supervised the experiments. Mohd. Shahir Shamsir and Talha Bin Emran supervised the research. Mohd. Shahir Shamsir and Talha Bin Emran acquisition the funding and visualized the investigation. All authors approved the final version of the manuscript.

## Availability of data and materials

All data generated or analysed during this study are included in this article and its supplementary data file.

## Declarations

### Ethics approval and consent to participate

Not applicable.

## Consent for publication

Not applicable.

## Supplementary Files

**Figure S1.** Secondary structure of template predicted by using GOR IV of network protein sequence analysis.

**Figure S2.** Secondary structure of model predicted by using GOR IV of network protein sequence analysis.

**Figure S3.** 3D Structure of luciferase-like monooxygenase from *P. meliae* (A0A0P9UTV8) as modeled using Alphafold Deepmind software. The head of the structure started from Methionine and ended of the amino acid is Alanine.

**Figure S4.** Surface structure of (**A)** Luciferase-like monooxygenase and (**B)** Template alkane monooxygenase.

**Figure S5.** The superimpose structure of template and model.

**Figure S6.** The surface image of the active site (blue) between model and template 3B9O. **Figure S7.** The surface image of active site (blue) template alkane monooxygenase from *Geobacillus thermodenitrificans* (strain NG80-2).

**Figure S8.** The probability of predicted active site (blue) of luciferase-like-monooxygenase from *Pseudomonas meliae*.

**Figure S9.** The structures between template and model has been highlighted by a different colour. The beta-strands are highlighted by blue colour, the loops are highlighted by grey colour and a shade of light brown highlights helices.

**Figure S10.** The strands of (**A)** Template alkane monooxygenase and (**B)** Model luciferase-like-monooxygenase.

**Figure S11.** The superimpose between model and template strands.

**Figure S12.** The loops of (**A)** Template alkane monooxygenase and (**B)** Model luciferase-like-monooxygenase.

**Figure S13.** The superimpose between model and template loops identify the similarities between model and template.

**Figure S14.** The helices between (**A)** Template alkane monooxygenase and (**B)** model luciferase-like-monooxygenase.

**Figure S15.** Helices superimpose between model luciferase-like monooxygenase and template alkane monooxygenase.

## Abbreviations

GRAVY: Grand average of hydropathicity
MSA: Multiple sequence alignment
ESPript: Easy Sequencing in PostScript
PDB: Protein data bank
3D: Three dimentional
2D: Two dimentional

